# MutPred2: inferring the molecular and phenotypic impact of amino acid variants

**DOI:** 10.1101/134981

**Authors:** Vikas Pejaver, Jorge Urresti, Jose Lugo-Martinez, Kymberleigh A. Pagel, Guan Ning Lin, Hyun-Jun Nam, Matthew Mort, David N. Cooper, Jonathan Sebat, Lilia M. Iakoucheva, Sean D. Mooney, Predrag Radivojac

**Affiliations:** Indiana University, Bloomington, Indiana, U.S.A.; University of California San Diego, La Jolla, California, U.S.A.; Cardiff University, Cardiff, U.K.; University of Washington, Seattle, Washington, U.S.A.

## Abstract

We introduce MutPred2, a tool that improves the prioritization of pathogenic amino acid substitutions, generates molecular mechanisms potentially causative of disease, and returns interpretable pathogenicity score distributions on individual genomes. While its prioritization performance is state-of-the-art, a novel and distinguishing feature of MutPred2 is the probabilistic modeling of variant impact on specific aspects of protein structure and function that can serve to guide experimental studies of phenotype-altering variants. We demonstrate the utility of MutPred2 in the identification of the structural and functional mutational signatures relevant to Mendelian disorders and the prioritization of *de novo* mutations associated with complex neurodevelopmental disorders. We then experimentally validate the functional impact of several variants identified in patients with such disorders. We argue that mechanism-driven studies of human inherited diseases have the potential to significantly accelerate the discovery of clinically actionable variants.

Availability: http://mutpred.mutdb.org/

## 1 Introduction

The discovery of pathogenic variants generally relies on a combination of family- and population-based sequencing efforts [1]. To assist genetic studies, particularly in characterizing rare variants and dissecting complex disease, machine learning methods have recently been developed to identify the signatures of pathogenicity and to predict the impact of variants of unknown significance [2, 3]. Although pathogenicity prediction methods have matured considerably over the past decade and are now routinely integrated into genomic pipelines, they continue to exhibit major shortcomings. Firstly, they remain inadequate to the task in exome-scale applications owing to a less than optimal balance of false positive and true positive detection rates [4, 5]. Secondly, they do not generate actionable hypotheses regarding the molecular consequences of these variants [6].

The functional impact of variants may lead to a wide range of molecular changes, even within a single protein, including disrupted stability and structure, disrupted macromolecular binding, ablation of post-translational modification (PTM) sites, among others (Fig. 1a). However, existing approaches generally provide little or no information about the potential mechanisms affected by mutations, or else simply map predicted pathogenic substitutions onto protein feature annotations (which are generally sparse) in public databases. These methods do not therefore explicitly model the type of change in local structure and function, and are fundamentally limited by the incomplete, incorrect and biased annotations in major databases [7–9].

**Figure 1:**
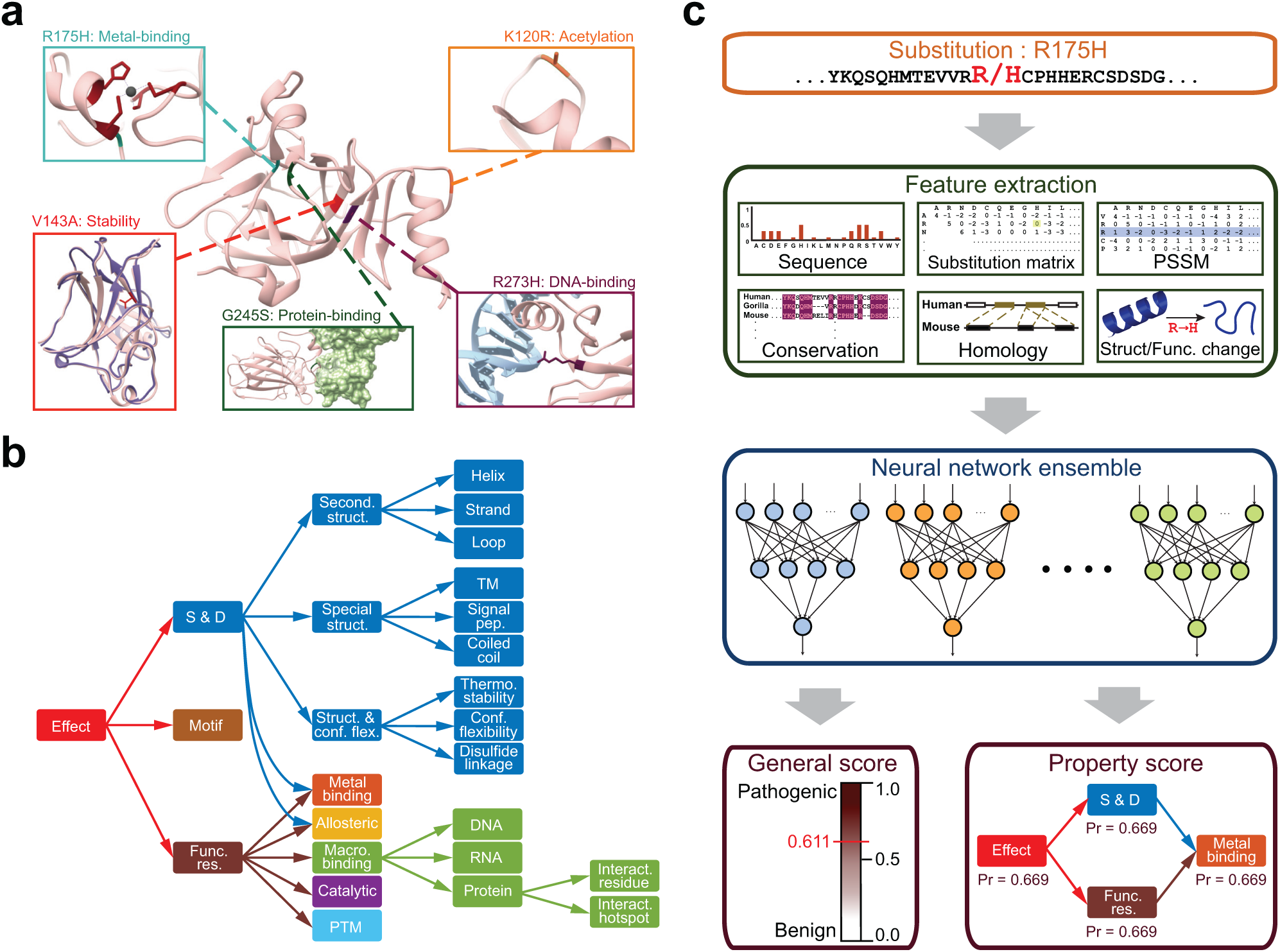
MutPred2 and the molecular consequences of amino acid substitutions. (a) The human tumor suppressor p53 as an illustration of the numerous possible effects of amino acid substitutions on protein structure and function. Protein Data Bank IDs for the structures shown are 1TUP, 1YCS, 2J1W and 2YBG. (b) The ontology constructed in this study to organize the possible structural and functional effects of amino acid substitutions. It is confined to the 53 properties included in MutPred2. (c) The MutPred2 workflow. For a given amino acid sequence and substitution, MutPred2 first extracts six categories of features from 25-residue fragments centered at the substitution position (except homolog counts, which are whole-protein features). Additionally, changes in structure and function due to the substitution are also modeled by running the original and mutated sequences through different sequence-based protein property predictors. Two scores are obtained for each property and these are combined to generate two additional scores quantifying the loss and gain of the property in question. All four scores are included as features. Next, all categories of features are presented to an ensemble of 30 neural networks trained to distinguish between pathogenic and benign variants. MutPred2 returns two outputs, the general score and the property score. The general score is obtained from the neural network ensemble and indicates the pathogenicity of the given variant. It ranges between zero and one, with a higher score indicating a greater propensity to be pathogenic. The property score is assigned to each of the 53 properties for the given variant and also ranges between zero and one. The latter score is the posterior probability of loss or gain (whichever is greater) of the given property due to the substitution. The higher the property score, the more likely that the molecular mechanism of the disease involves the alteration of the property.

To address these challenges, we have extended our existing and widely used machine learning approach, MutPred [10], by developing a novel and statistically rigorous approach, MutPred2. This new algorithm quantifies the pathogenicity of amino acid substitutions and describes how they affect the phenotype by modeling a broad repertoire of structural and functional alterations from amino acid sequence.

MutPred2 compares favorably with the existing tools recommended in the ACMG Standards & Guidelines [11] on a stringent independent test set. More importantly, by applying this methodology we estimate the fraction of deleterious missense variants in a personal genome and identify molecular signatures associated with a data set of Mendelian disease variants and a data set of *de novo* mutations found in individuals diagnosed with neurodevelopmental disorders. Finally, we prioritize several high-scoring variants from this data set and experimentally validate their functional roles. Our results suggest new molecular targets and mechanisms impacted by multiple mutations across neurodevelopmental disorders. More broadly, this study demonstrates the power of a novel mechanism-driven approach to studying human phenotypes.

## 2 Results

MutPred2 is a machine learning-based method and software package that integrates genetic and molecular data to reason probabilistically about the pathogenicity of amino acid substitutions. This is achieved by providing (1) a general pathogenicity prediction, and (2) a ranked list of specific molecular alterations potentially affecting the phenotype. MutPred2 is a sequence-based model that utilizes methodology predicated upon recent machine learning advances in training from positive-unlabeled data, and which incorporates estimation of prior and posterior probabilities [12, 13]. These estimates in turn facilitate the interpretation of pathogenicity and molecular alteration scores as well as provide a framework to rigorously rank the underlying mechanisms [13]. Currently, MutPred2 models a broad range of structural and functional properties, including secondary structure, signal peptide and transmembrane topology, catalytic activity, macromolecular binding, PTMs, metal-binding and allostery.

### 2.1 Challenges for the development of next-generation interpretation tools

To develop models for the mathematically sound inference of molecular mechanisms of disease, several statistical and computational challenges must be addressed. First, it is necessary to integrate disparate molecular and genetic data to develop models that have similar yet meaningful score interpretations [6]. Second, prediction software tools generally vary not only in their feature representation and prediction algorithms, but also in their implementations, dependencies, and system requirements, which collectively hinder the development of a robust framework that seamlessly incorporates multiple models. Third, structural and functional properties occur with unequal prior probabilities, requiring sophisticated modeling for ranking the properties affected by a substitution. Finally, although these property predictors are typically developed independently of one another, they are interrelated; i.e., a single substitution may affect more than one property. This places the burden of interpretation upon the user and can be overwhelming when multiple properties are considered simultaneously.

To address the first two challenges, we developed sequence-based predictors for over fifty structural and functional protein properties (Supplementary Tables 1-3). All predictors, with minor exceptions, were trained with a common feature set, subjected to the same evaluation protocols, designed to output scores between 0 and 1, and implemented within the positive-unlabeled machine learning framework (Supplementary Materials). The areas under the ROC curves (AUCs) of these predictors are generally high (Supplementary Table 4). To address the third challenge, we estimated the prior probability for each property [13, 14] and used it to transform raw prediction scores to posterior probabilities to facilitate direct comparisons with other properties [13] (Supplementary Table 5). To address the fourth challenge, we constructed a custom ontology of molecular alterations by grouping the properties into broader categories to capture the inherent relationships between the properties (Fig. 1b). This was carried out manually by combining our current understanding of protein structure and function with the Variation Ontology as a template [15]. Our primary goal was to organize the space of molecular consequences so as to achieve user-friendly interpretation (Fig. 1c).

The MutPred2 pathogenicity model was trained on a set of 53,180 pathogenic and 206,946 unlabeled (putatively neutral) variants obtained from the Human Gene Mutation Database (HGMD) [16], SwissVar [17], dbSNP [18] and inter-species pairwise alignments. The model is a bagged ensemble of feed-forward neural networks [19], each trained on a balanced subset of pathogenic and unlabeled variants. The final prediction score is the average of the scores from all networks and ranges between 0 and 1; higher scores reflect a higher probability of pathogenicity. MutPred2’s models for inferring molecular mechanisms were similarly trained from a variety of molecular data sets (Supplementary Materials), thereby ensuring effective integration of genetic and molecular data.

### 2.2 Evaluation of predictor performance

The choice of the training set is critical in machine learning. A common practice in pathogenicity prediction involves the exclusion of rare variants from the unlabeled set to minimize biases arising from potentially undiscovered pathogenic variants; additional filtering based on specific types of data source may also be performed. To investigate the effects of various filtering criteria in training sets on classification performance and to select the most appropriate training set, different combinations of training and test sets were evaluated in an all-against-all performance assessment (Supplementary Table 6). We found that filtering the training set is beneficial only when the test set is also filtered using the same criteria. Furthermore, using the entire unfiltered training set resulted in comparable or better performance in most cases, consistent with recent theoretical results justifying training from positive-unlabeled data [12, 13]. Therefore, we chose not to perform any filtering in subsequent steps with the reasoning that bias introduced through different filtering schemes is more detrimental than random noise [12, 13].

Using a per-protein 10-fold cross-validation, the area under the ROC curve (AUC) was estimated at 87.7% (Fig. 2a). The training data, however, contains class-label noise; i.e., the set of disease variants may contain mutations incorrectly labeled as pathogenic and the set of unlabeled variants is by definition also a mixture of pathogenic and benign variants. We estimate the proportion of noisy positive variants to be 2.8% and the proportion of unlabeled variants that are pathogenic to be 5.8% (Section 2.5; Supplementary Materials). These results allow us to provide a corrected estimate of the AUC of 91.3% [20].

Consistent with previous studies, conservation-based features were the most discriminative [4, 21, 22] (Supplementary Table 7). MutPred2 relies on precomputed databases of multiple sequence alignments and conservation scores to calculate these features. In cases where the input substitutions come from novel protein sequences or alternate isoforms, these data may often be unavailable, prompting tools to avoid assigning predictions. To ensure that every input variant has a prediction, MutPred2 provides an option of predicting conservation-based features from sequence and PSI-BLAST position-specific scoring matrices (PSSMs) in cases when these features are unavailable. Although predicted conservation features only moderately correlate with actual values (Supplementary Fig. 1), models that include them performed two percentage points better than those that did not use any conservation features (Fig. 2a).

**Figure 2:**
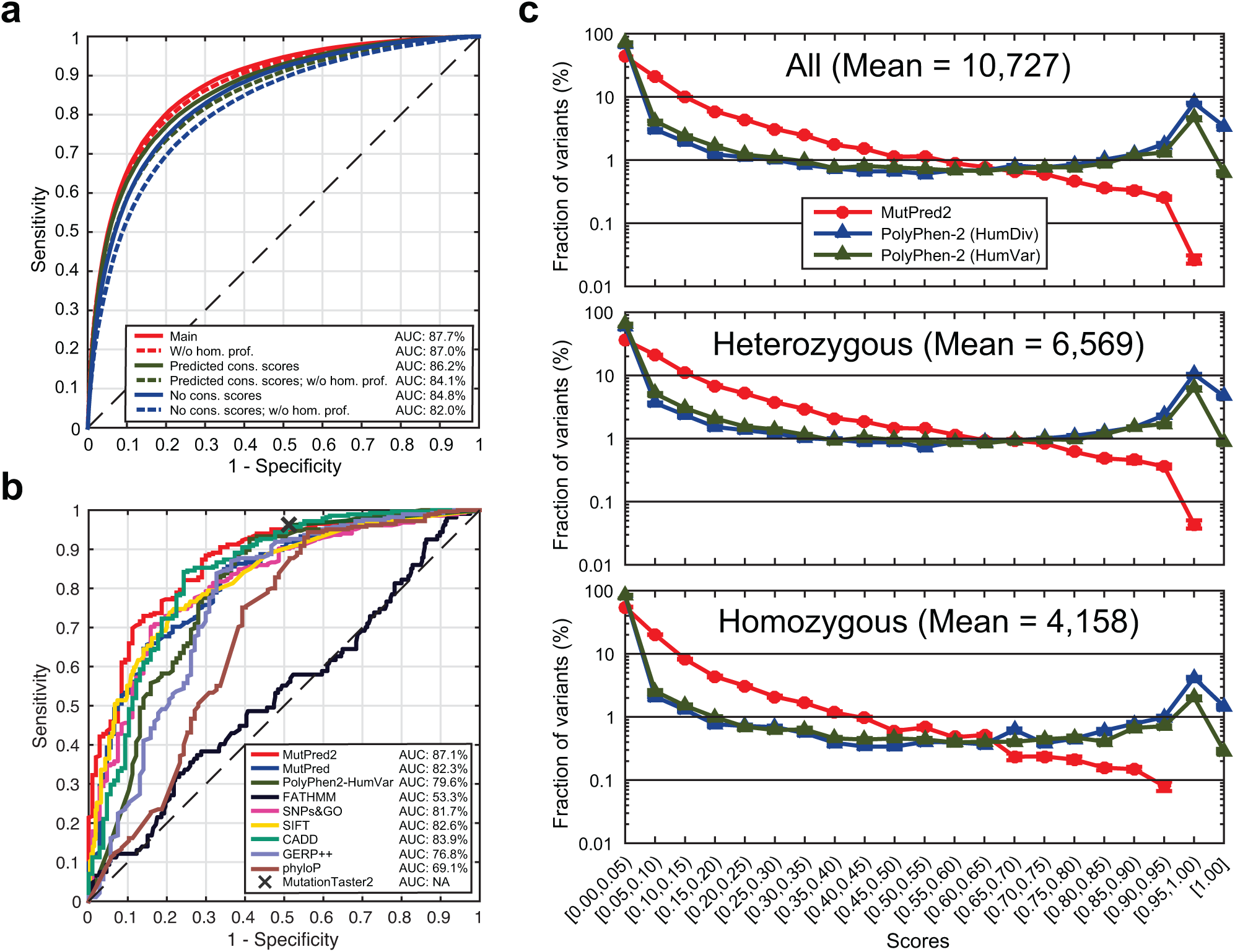
Performance and interpretability of MutPred2. (a) ROC curves obtained through 10-fold cross-validation on the MutPred2 training set. The main model represents MutPred2 in the default setting (with real conservation scores and homolog count profiles). All lines are paired with the solid line representing the model with homolog count profiles and the dashed line representing the model without the profiles. (b) ROC curves on an independent test set, obtained from ClinVar and SwissVar by letting the data accumulate in these databases for three years. MutationTaster2 only returns a value of zero or one and therefore its performance is plotted as a single point (“X”). Since some tools could not assign scores to all variants, results from the subset of the variants (285 pathogenic and 107 benign) that are covered by all methods are shown. Detailed performance measures on this subset and a less stringent set (filtered at 80% sequence identity) are shown in Supplementary Tables 8-9. (c) Mean score distributions for MutPred2, PolyPhen-2 HumDiv and PolyPhen-2 HumVar applied to 10 randomly selected exomes from the 1000 Genomes Project. All heterozygous and homozygous variants were plotted in separate panels. The “Mean” represents the average number of variants found in an individual for the given category.

Previous studies reported conflicting results on the association between duplicated genes and their involvement in disease [23, 24]. To investigate this, we created features that, for a given protein, enumerate homologous proteins from human and mouse at various levels of sequence similarity (50% or greater; Supplementary Materials). We observed that excluding these homology profiles did not drastically affect performance when true conservation features were available, but resulted in a decrease of two percentage points in all other cases (Fig. 2a). This outcome supports the evidence for compensatory mechanisms in a variety of gene families [24].

We then evaluated MutPred2 against MutPred on its original training set under the same evaluation protocol. We found that MutPred2 had a similar AUC value as before (88.0%), outperforming the original MutPred approach by about 5 percentage points (Supplementary Fig. 2). Additional experiments are described in Supplementary Materials.

### 2.3 Evaluation against external tools

We next compared the performance of the default MutPred2 model with several state-ofthe-art methods recommended in the ACMG Standards & Guidelines on the interpretation of sequence variants [11]; specifically, CADD [25], FATHMM [26], GERP++ [27], MutationTaster2 [28], MutPred [10], PhyloP [29], PolyPhen-2 [22], SIFT [21], and SNPs&GO [30]. This analysis was carried out on an independent data set compiled from ClinVar [31] and SwissVar. We note that such comparisons are limited by differences in motivating goals, problem formulations, training data and information used to make predictions, and are best addressed through community-wide challenges such as the Critical Assessment of Genome Interpretation; CAGI (http://genomeinterpretation.org). For a fair and rigorous comparison, we minimized potential biases by including only variants that were not in the training sets (where known) of the methods listed above. The analyses were restricted to a non-redundant set of variants by comparing 25-residue fragments centered at the variant position to all such fragments in these training sets and filtering out those that shared more than 50% sequence identity. This resulted in a highly stringent data set of 343 pathogenic mutations and 137 non-pathogenic mutations.

We found that MutPred2 performed substantially better than FATHMM, GERP++, MutationTaster2, PhyloP and PolyPhen-2 in terms of AUC (Fig. 2b, Supplementary Table 8). The remaining methods resulted in AUCs of at least 80% and ROC curves that grouped together. However, MutPred2 still had the highest AUC (87.1%), largely attributable to its high sensitivities at lower false positive rates (FPRs). This is especially relevant when considering the second-best performing tool, CADD. Despite the possibility that some of these variants may have been present in its training set and the fact that its AUC value was very close to that of MutPred2, CADD was more sensitive only when the FPR was high (between 20% and 30%). Interestingly, contrary to results from previous studies [32, 33], the next best-performing method on this independent data set was SIFT. This is possibly due to the use of a newer version of the software.

Given that most methods considered here do not return predictions for some fraction of the independent data set, the small data set size limits the interpretability of these findings. To mitigate this, we relaxed the fragment sequence identity threshold to 80% and expanded the independent data set to include 700 pathogenic mutations and 282 non-pathogenic mutations. Although individual performance values changed, the general trends remained unaffected, with the exception of SIFT’s reduced performance (Supplementary Table 9).

### 2.4 Interpretability of prediction scores

The interpretability of prediction scores is a generally underappreciated aspect of pathogenicity prediction. From this perspective, it is desirable that predicted pathogenic variants show sufficient dispersion of scores so that they can be further grouped into meaningful bins for human interpretation. It is also desirable that the scaling of scores is linear. For instance, for two variants with scores of 0.9 and 0.7 respectively, one should be able to infer that both are pathogenic but that the variant scored 0.9 is much more likely to be pathogenic. However, for variants with scores 0.82 and 0.80, such an interpretation would be problematic. Intuitively, one should also be able to interpret the difference between 0.9 and 0.7 in a manner similar to that between 0.7 and 0.5; i.e., quantitative differences should reflect qualitative differences.

To visualize this, we applied MutPred2 and PolyPhen-2 to missense variants from 10 randomly selected presumably healthy individuals from the 1000 Genomes Project [34] and plotted the resulting score distributions (Fig. 2c). We found that, while pathogenic and benign predictions for PolyPhen-2 tended to peak at the tails of the distribution, MutPred2 outputs a generally decreasing score distribution. We believe that MutPred2’s distribution is better suited for the interpretation of personalized genome-scale data than the bimodal PolyPhen-2 distribution, as it allows for the interpretation of scores as crude probabilities.

The shapes of the distributions of the two methods also suggest a difference in false positive prediction rates when applied to whole genomes. Although these two distributions are not directly comparable, we considered a high threshold range of at least 0.9 to mitigate any effects arising from differences in sensitivity-specificity values at lower thresholds. At this cutoff, MutPred2 predicts 0.3% of all amino acid substitutions to be pathogenic, which is an order of magnitude lower than the 6.6% predicted to be pathogenic by PolyPhen-2 (Fig. 2c), even when considering the better-suited HumVar model in PolyPhen-2. We initially thought that such a high number of predicted pathogenic variants could potentially be attributable to zygosity; however, these trends also hold when considering only homozygous or heterozygous variants. MutPred2 still predicted far fewer homozygous variants to be pathogenic than PolyPhen-2 (9 vs. 322, Fig. 2c). We also investigated whether these distributions were impacted by minor allele frequencies (MAFs). We found that MutPred2 scores were better anti-correlated with MAFs (Supplementary Fig. 3), further suggesting that MutPred2 scores align better with theoretical expectations of the allele frequencies of slightly deleterious variants.

### 2.5 Estimating proportions of pathogenic missense variants

The set of unlabeled substitutions in the MutPred2 training data comprises both benign and pathogenic variants that have not yet been characterized or annotated as such. This is also the case for the set of substitutions labeled as pathogenic as a consequence of possible errors and misannotations in our positive set. To quantify these proportions in our training set, we generated MutPred2 score distributions on these sets and applied the AlphaMax algorithm [13]. On our training set, we estimate that 5.8% of the unlabeled variants may indeed be pathogenic and that 2.8% of the pathogenic variants may actually not be associated with disease (Supplementary Fig. 4). Using these class prior estimates as proxies for those occurring in nature, we then estimated the average proportion of pathogenic missense variants over the genomes of the 10 individuals from the 1000 Genomes Project. We found that, on average, up to 1.3% of all missense variants in a (presumably) healthy genome are pathogenic. This fraction was greater for heterozygous variants (1.7%) than for homozygous variants (0.6%).

### 2.6 Enriched molecular mechanisms in Mendelian diseases

We sought to identify preferentially disrupted mechanisms in the MutPred2 training set by calculating the enrichment of the increase (i.e., gain) and decrease (i.e., loss) of local structural and functional properties in the set of disease mutations relative to the unlabeled variants (FPR of 1%). We recapitulated previous observations that protein structural changes are common mechanisms of Mendelian disorders (Fig. 3) [35, 36]. However, we also observed enrichment for macromolecular binding sites, in agreement with recent work on protein-DNA and protein-protein interactions (PPIs) [37], as well as several PTM types. We observed depletion for properties associated with flexible and disordered regions, potentially owing to the enrichment of enzymes involved in metabolic processes [38]. Additionally, we found that substitutions affecting residues involved in metal binding and allosteric regulation were also enriched in the set of Mendelian disease mutations.

**Figure 3:**
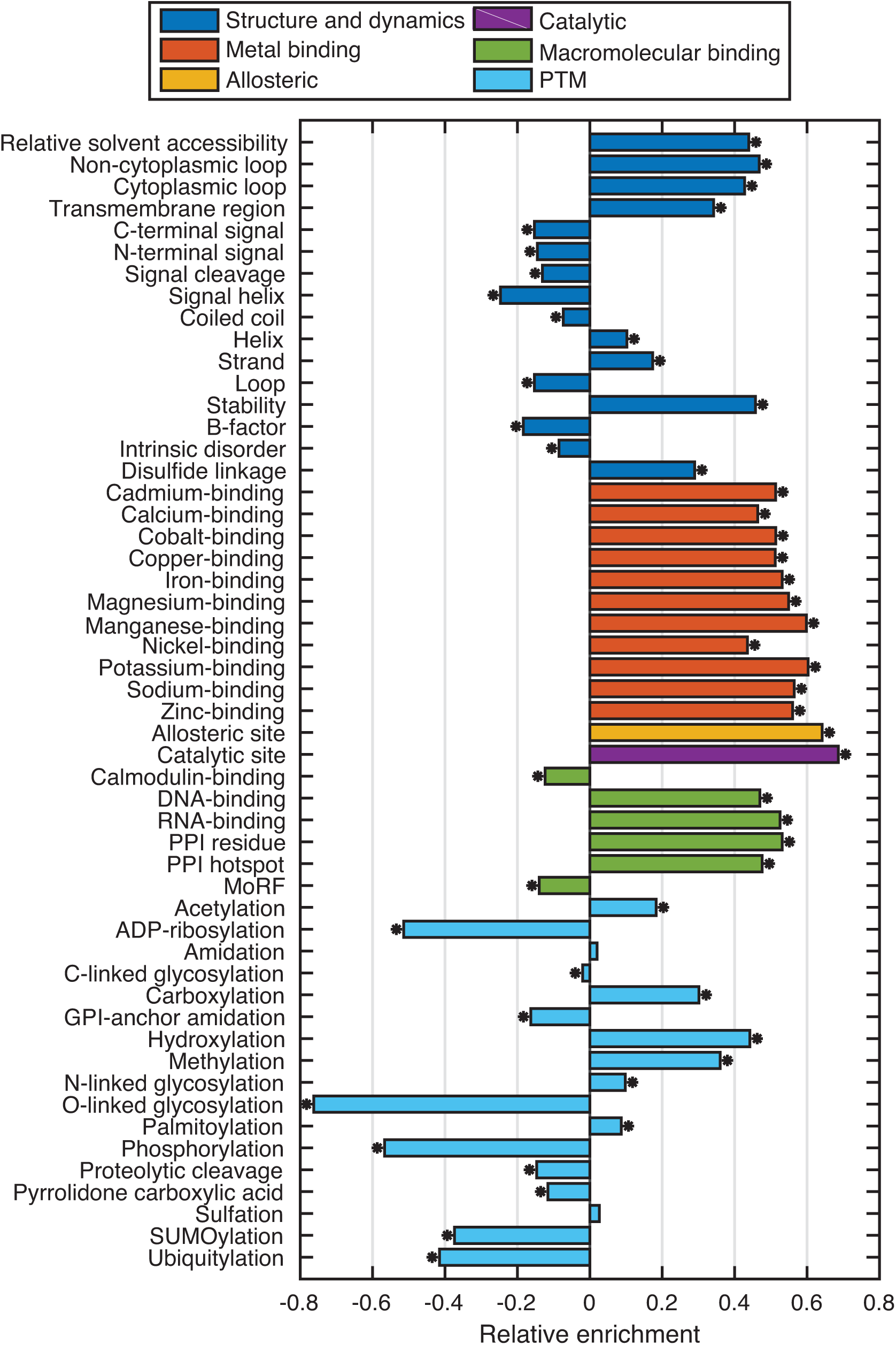
Significantly enriched and depleted pathogenic mechanisms in the inherited disease mutation data set, as predicted by MutPred2. Losses and gains are plotted together by considering the maximal effect for a given mutation position. An ‘*’ indicates significance at the 0.05 level, with Benjamini-Hochberg correction.

Metals, unlike PTMs, freely form coordinate bonds without enzyme involvement and are perhaps more ubiquitous in nature. One metal ion can also be in competition with another at one or more sites in a protein due to their similar chemical properties [39]. In terms of allosteric regulation, to the best of our knowledge, a sequence-based predictor of allosteric residues does not exist, and only one structure-based study has systematically investigated the role of allosteric regulation in monogenic diseases [35]. Contrary to the findings of that study, we predict that the disruption of allosteric sites is an active mechanism in such diseases.

Allosteric regulation and metal binding are treated as both structural and functional properties in our ontology (Fig. 1b). This is supported by the fact that metals are known to play important roles in stabilizing both protein structure and macromolecular interactions [39]. Further details of the enrichment analysis are provided in Supplementary Materials (Supplementary Fig. 5).

### 2.7 Structural and functional signatures of *de novo* missense mutations in neurodevelopmental disorders

Neurodevelopmental disorders have a strong genetic component [40, 41]. Recent wholegenome and whole-exome sequencing in neurodevelopmental diseases has identified thousands of *de novo* mutations in patients with such phenotypes. However, distinguishing between pathogenic and benign *de novo* mutations remains challenging. We applied MutPred2 to a data set of 4,324 *de novo* missense mutations obtained through the exome sequencing of families impacted by four different disorders (autism spectrum disorder, intellectual disability, schizophrenia and epileptic encephalopathy), and 1,316 *de novo* missense mutations from the healthy siblings of the patients from these families as a control (Methods).

We first examined whether pathogenicity scores alone were sufficient to distinguish between cases and controls. We found that MutPred2 predicted significantly higher proportions of pathogenic mutations in the case set than in the control set at 10% and 5% FPR score thresholds with odds ratios of 1.44 and 1.56, respectively (Fig. 4a). Statistically significant odds ratios exceeding 1.22 were observed starting at a score threshold of 0.45 (Supplementary Fig. 6). Given the fact that the overall mutational load for *de novo* missense mutations is similar in the cases and the healthy controls [42, 43], the higher fraction of predicted pathogenic missense mutations in the cases suggests good discriminative ability of MutPred2. Low odds ratios are not unexpected, as missense variants are likely to be less disruptive to protein structure and function than loss-of-function (stop, frameshifting indel, splice site) variants, for which a 2-fold enrichment in cases vs. controls has been previously observed for autism [42].

**Figure 4:**
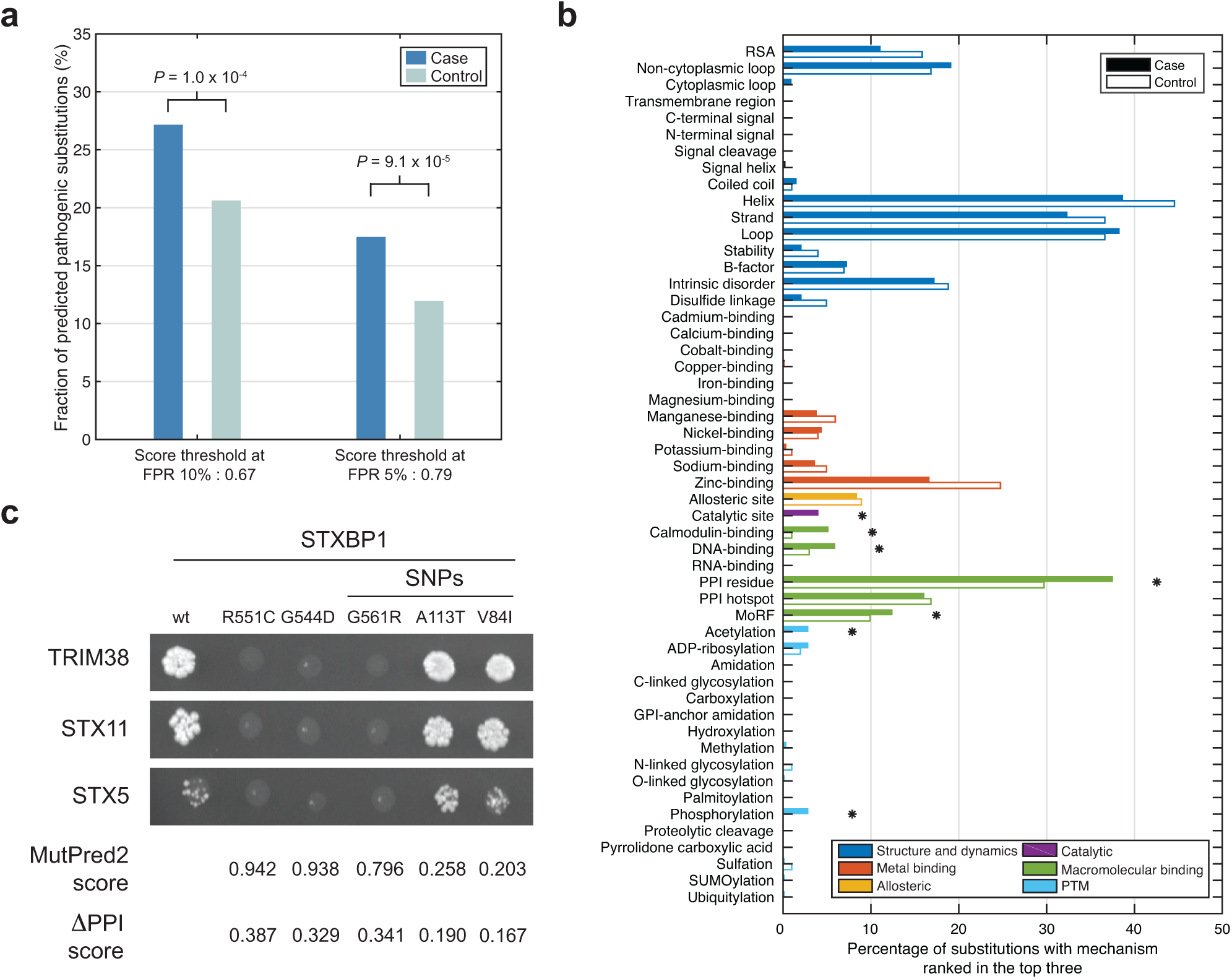
Summary of MutPred2 predictions on *de novo* missense mutations from four neurodevelopmental disorders. (a) Proportions of case and control mutations predicted to be pathogenic by MutPred2 at thresholds corresponding to false positive rates (FPR) of 10% and 5% respectively. P-values were computed using Fisher’s exact test. Odds ratios and P-values for other thresholds are shown in Supplementary Fig. 6. (b) Enrichment of structural and functional signatures of case mutations versus the control group. Only those mutations considered to be pathogenic at the 5% FPR threshold were included in this analysis. Properties are grouped based upon their broader classes as described in the ontology (Fig. 1b). Statistical significance was assigned at *α* = 0.05 using a one-sided binomial test with Benjamini-Hochberg correction. (c) Representative images of 3AT selection plates with the interaction profiles of STXBP1 against TRIM38, STX11 and STX5. Pathogenicity scores and probability of alteration of PPIs corresponding to each mutation are shown. The PPI alteration probability of greater than 0.5 (1 0.5) = 0.25 is considered to be high-scoring.

Next, we examined whether mutations in neurodevelopmental disorders preferentially alter specific protein structural and functional properties. We asked which molecular mechanisms were frequently ranked among the top three in the set of predicted pathogenic mutations (at the 5% FPR threshold) from the case and control sets (Supplementary Table 10). In contrast to the set of Mendelian disorders, we observed the statistically significant enrichment of the majority of macromolecular binding features (e.g., calmodulin binding, DNA binding, protein binding), catalytic sites, and two types of PTMs (acetylation, phosphorylation) among the case mutations (Fig. 4b). The most significant enrichment was observed for the PPI residue feature, in agreement with previous studies demonstrating the loss of protein-protein interactions as a result of mutations associated with human Mendelian diseases [44–46]. At the same time, well-defined structure and metal binding were not significantly enriched among case mutations, despite high proportions of secondary structure elements (helix, strand and loop) being affected by the mutations in both cases and controls (Fig. 4b).

### 2.8 Experimental validation of the functional impact of *de novo* missense mutations

High pathogenicity scores for a given mutation provide hypotheses of disruption of protein structure or function that could lead to a disease. We used the yeast-two-hybrid (Y2H) system to test the impact of our high-scoring mutations on protein-protein interactions with corresponding binding partners as previously described [44, 45].

We selected three genes with *de novo* missense mutations: *STXBP1*, *ZBTB18* and *DNMT3A*, and introduced the high-scoring mutations into the open reading frame (ORF) clones of these genes. We then tested both wild-type and mutant variants for interactions with protein partners that were available in the human hORFeome collection [47] (Methods). We also tested common single nucleotide polymorphisms (SNPs) from dbSNP adjacent to the *de novo* mutations as experimental controls.

Two mutations, R551C [48] and G544D [49], and one SNP (G561R) in the *STXBP1* gene disrupted interactions between STXBP1 and three protein partners (TRIM38, STX11 and STX5), whereas two other SNPs (A113T and V84I) did not alter interaction patterns compared to the wild-type protein (Fig. 4c). These results are in good agreement with the MutPred2 predictions that assigned high scores to both R551C (score 0.942) and G544D (score 0.938) and also to the G561R (score 0.796) SNP. Interestingly, although the G561R variant is present in dbSNP and has unknown clinical significance, it is a rare variant in the ExAC database [50] (MAF = 9.2 · 10^*-*6^). STXBP1 is a syntaxin-binding protein that plays a role in the release of neurotransmitters via regulation of syntaxin, a transmembrane attachment protein receptor. A recent study demonstrated that heterozygous loss of *STXBP1* in human neurons lowers the level of its protein product together with its binding partner syntaxin-1, emphasizing the importance of this protein-protein interaction [51]. STXBP1 is highly expressed in the brain, and other mutations in *STXBP1* are associated with early infantile epileptic encephalopathy [49], intellectual disability [52] and developmental delay [53]. Our results suggest that the underlying mechanism of the mutations tested here could be attributable to loss of binding to protein partners.

We then examined the effect of the R486G mutation [52] and three SNPs (G416R, A502T, and T507A) on the binding of ZBTB18 with two partners: CTBP1 and CTBP2 (Supplementary Fig. 7a). Although the experimental results for R486G (score 0.932), G416R (score 0.668) and T507A (score 0.069) were in agreement with predictions, the MutPred2 score for A502T (score 0.208) did not match the observed loss of binding (Supplementary Fig. 7a). A502T may therefore be an undiscovered pathogenic variant missed by MutPred2 or alternatively it is a PPI-altering variant that does not lead to a disease phenotype. Finally, we tested the R635W mutation [54] in the *DNMT3A* gene (Supplementary Fig. 7b). The prediction result for R635W (score 0.886) agreed with experimental observation of the loss of interaction with the TCL1A partner.

Overall, MutPred2 predictions agreed well with experimental observations. It is, however, important to interpret these results with caution, because the loss of PPIs could also result from the loss of stability, aberrant folding, increase in degradation, etc.

## 3 Discussion

An individual person’s genome contains about 10,000 amino-acid altering variants when compared to the reference genome [34]. The first step in connecting this information with phenotype, and particularly with disease, involves prioritizing variants that affect protein structure and function. The current generation of pathogenicity prediction methods has enabled the reduction of a large number of variants to hundreds of possible candidates. However, these numbers remain prohibitively large for subsequent experimental characterization, even considering high-throughput studies *in vitro* or new CRISPR/Cas9 technologies. To address this challenge, we developed MutPred2, a tool for the inference of the structural, functional and phenotypic consequences of sequence variants. By modeling the effects of variants on local protein structure and function, MutPred2 improves pathogenicity prediction and assigns putative molecular alterations using a novel ranking approach. This additional information can be used to accelerate experimental validation. The assignment of specific molecular impact also allows one to quantify molecular signatures within different data sets; e.g., classes of disease, a specific disease, a healthy population, a subpopulation, among others.

### 3.1 Larger and more heterogeneous training sets result in better models

Previous studies have developed predictors trained on the data sets that were filtered based on minor allele frequencies (MAFs) and/or source of data [22, 55]. Our results suggest that such filtering is beneficial only when it is directly relevant to the prediction task at hand (Supplementary Table 6); in fact, we observe that the model trained on the entire data set without filtering performs well across all prediction tasks, thereby reducing the need for specialized models. We attribute this good generalization performance to the availability of more data (the data set size decreases drastically when frequency-based filtering is applied) and reduced ascertainment bias.

### 3.2 An individual’s genome contains about 1.3% pathogenic missense variants

There has been debate on the fraction of missense variants in an individual’s genome that contributes to disease. Estimates for the proportion of missense variants in a genome that have deleterious effects on protein function, and hence phenotype, have ranged between 10% and 25%, depending on the operationalization of pathogenicity [56–58]. However, other works have suggested that this proportion could be much lower [59–61]. In general, the calculation of this fraction on real data has been confounded by the use of small and biased data sets, the accuracy of the underlying pathogenicity prediction method, the need to make estimates at a predetermined false positive rate, the limitations of simplifying assumptions on the parameters of the relevant theoretical framework, and differences in terminology.

The combined use of MutPred2 and AlphaMax [13] on exome-scale data allowed us to address these issues in a rigorous manner, with few assumptions. We established that, on average, about 100 heterozygous and 25 homozygous variants in an individual may cause disease, on par with estimates derived using disease-causing variants from HGMD [61]. Although these numbers are large enough to yield disease phenotypes, our estimates do not account for epistatic interactions such as compensatory variants [62]. However, our estimates are the lowest among those derived directly from data generated thus far and support the views of early studies [60, 62].

From a practical perspective, the extent of noise in current training sets for pathogenicity prediction is also important. Our results suggest that incorrectly labeled pathogenic variants constitute a small fraction of our training set and may not seriously impact predictive performance. Our estimates also suggest that around 10,000 pathogenic amino acid substitutions in dbSNP and UniProt may currently be unannotated. We believe this result is reasonable, considering that we did not filter out rare variants in our unlabeled set. However, it is important to note that our work does not address the issues of bias in current training sets [13, 63].

### 3.3 Some *de novo* missense mutations implicated in neurodevelopmental diseases disrupt protein interactions

Genes with *de novo* mutations are strongly associated with neurodevelopmental disorders. Several studies have demonstrated that the protein products of these genes physically interact and form tightly connected protein interaction networks [64–66]. However, how specific mutations impact interactions between proteins and which mutations are pathogenic remains an open question.

MutPred2 predicts more pathogenic *de novo* missense mutations in cases than in controls. This is particularly remarkable given that brain-relevant information that could increase predictor’s performance for this type of disease was not exploited. In fact, the only filtering step involved the removal of genes present in both the case and control sets; even without this step, statistically significant odds ratios greater than one were obtained (not shown). Although the log odds are moderate, each missense variant can now be associated with potential molecular alterations that should prompt further investigation (Supplementary Table 10). For example, the L834P mutation in CHD8 identified in a patient with autism [67] is predicted to disrupt a catalytic site along with the allosteric site and PPI residue. The mutation K842R nearby has been shown to abolish the ATPase activity of CHD8 [68], which is consistent with the MutPred2 prediction of catalytic site disruption by an adjacent L834P mutation. Likewise, the M2679T mutation in the RYR3 calcium channel identified in an autism patient [69] is predicted to disrupt calmodulin binding along with a loss of helical propensity. By similarity with other ryanodine receptors, RYR3 probably binds calmodulin at its C-terminus, and the prediction of the loss of calmodulin binding due to the M2679T C-terminal mutation concurs with the known function of this protein. Thus, MutPred2 predictions offer viable biological hypotheses that can be tested in the laboratory to improve our understanding of disease mechanisms.

### 3.4 Molecular mechanisms as drivers in disease studies

Traditionally, researchers have adopted a top-down or disease-driven approach, where one starts with specific phenotypes and works one’s way down towards causes at the molecular level. MutPred2 enables the adoption of a bottom-up or mechanism-driven approach towards understanding genetic disease. In this approach, one can envision experts specializing in molecular mechanisms studying germline and somatic variants across different diseases and providing functional insights that can subsequently lead to hypotheses at the phenotypic level [70, 71]. We loosely refer to this mechanism-driven approach as disease-agnostic because the study and validation of impactful variants is determined by molecular mechanisms one is equipped to study, not necessarily the high-level phenotype. By grouping disease classes together based on frequently affected molecular mechanisms in current data sets, one can consider the prospect of identifying common targets and repurposing drugs from one class of disease to its neighboring disease in this new space. For example, MutPred2 predicted a close relationship between the endocrine and immune systems at the molecular level (Supplementary Fig. 5). This agrees with observations related to the interactions between the two systems during ontogeny [72].

## 4 Methods

An input to MutPred2 is an amino acid sequence *s*; i.e., a wild-type (wt) protein sequence, and an amino acid substitution *XiY*, where *X*, the *i*-th amino acid in *s*, is replaced by *Y*. We refer to the mutated (mt) sequence as *s*_*XiY*_. The output of MutPred2 consists of a pathogenicity score, a number from [0, 1], and a list of molecular mechanisms, each with its own score, that may be impacted by *XiY*. A pathogenicity score of 1 indicates near-certainty that the variant is pathogenic, whereas a score of 0 indicates near-certainty that the variant is benign. In the next several sections we discuss data sets, data representation and training of MutPred2. The details regarding classification models used to assess specific functional impacts are provided in Supplementary Materials.

### 4.1 Data sets

A data set of pathogenic amino acid substitutions was created by integrating HGMD [16] (June 2013; “DM”-annotated substitutions only), Swiss-Prot (release 2012 09 through SwissVar [17]) and dbSNP [18] (build 137). The set of unlabeled (putatively neutral) substitutions was compiled from Swiss-Prot and dbSNP, and then supplemented with additional variants in a way similar to the HumDiv training set for PolyPhen-2 [22]. Specifically, for every human protein, pairwise alignments to other mammalian proteins were first extracted from a 46-species multiple sequence alignment, obtained from the UCSC Genome Browser [73]. Only those alignments where the two sequences shared at least 99% sequence identity were considered and positions where a residue in the non-human sequence was replaced by a different one in the human sequence were identified.

### 4.2 Data representation and training

Given a sequence *s* and variant *XiY*, we extracted 1,345 (including 20 optional) features. These features are subcategorized into six groups: (1) sequence-based features, (2) substitution-based features, (3) position-specific scoring matrix-based features, (4) conservation-based features, (5) homolog profiles (optional due to time necessary to compute), and (6) changes in predicted structural and functional properties. A detailed list of features and how they were extracted and encoded is provided in Supplementary Materials. Feature selection using a two-sample t-test was performed and only those features that returned P-values less than 0.01 were retained. To remove (near) co-linear features, z-score normalization and principal component analysis (PCA) were performed on the selected features, with the retained variance set to 99%. An ensemble of 30 feed-forward neural networks was then trained on the resulting feature matrix. Each network consisted of a single hidden layer with four neurons and a single output neuron (the hyperbolic tangent activation function was used in both layers). A bootstrap aggregating (bagging) approach was adopted for training, where each network was trained on a balanced random sample (with replacement) of the original training set. To determine the number of iterations required for training, 25% of the training data were retained as a validation set. The final model was trained using the resilient propagation algorithm [74] and stopped when, either this optimal number of iterations was reached, or 1000 epochs were completed, or 500 validation checks were reached. Prediction scores were then calculated as the mean of all 30 scores.

### 4.3 Inferring molecular mechanisms of pathogenicity

The local effects of a variant on predicted structural and functional properties were modeled and utilized, both as features and for the assignment of putative molecular mechanisms. First, over 50 protein property predictors were developed within a unified positive-unlabeled learning framework (Supplementary Materials). The wt sequence *s* was first provided to these predictors to score the substitution site *i* and ±5 adjacent positions. The amino acid substitution was then introduced into the sequence in silico and the mt sequence *s*_*XiY*_ provided to all property predictors. The probabilities of changes in structural and functional propensities, given the substitution *XiY*, were calculated from the property predictors as follows

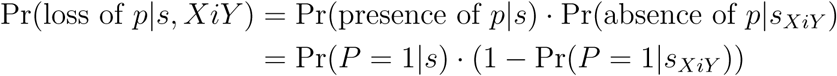

and

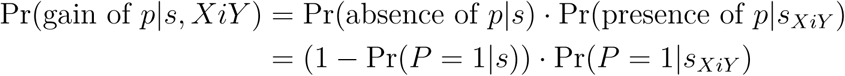

where *P* is the random variable indicating presence (1) or absence (0) of property *p*. In the above equations, the wild-type residue at the *i*-th position of the protein is *X* and the replacement amino acid is *Y*. Appropriate transformations were applied to ensure that the property predictors accurately approximate posterior distributions (Supplementary Materials). The posterior probability from the predictor for property *p* for the wild-type sequence can be interpreted as Pr(*P* = 1 | *s*), and the posterior probability for the substituted sequence can be interpreted as Pr(*P* = 1 | *s*_*XiY*_).

The property score was interpreted as the posterior probability of loss or gain, whichever was greater. Naively, if the wild-type posterior is 0.5 and the mutant posterior is 0.5 (i.e., no effect upon substitution), then the loss and gain probabilities will be 0.25, which we treated as a baseline threshold to implicate the property as a molecular mechanism in disease. It is important to note that the terms “loss” and “gain” are more appropriately interpreted as decreased and increased propensities for a certain property, respectively. Furthermore, in the case of properties that can be affected in both directions due to a single-residue change, interpretation becomes complicated. For instance, a substitution can increase a protein’s propensity to bind one protein partner but decrease its propensity for another. For simplicity, the term “altered” is reported in MutPred2 predictions for such properties along with the maximum of the loss and gain score. In addition to posterior probabilities of loss and gain, we also provide empirical P-values that the observed loss/gain score is as high or higher than the score randomly generated from the distribution of putatively neutral substitutions. The lower this P-value, the more likely that the predicted property is involved in pathogenicity, under the assumption that non-pathogenic variants do not affect protein structure and/or function.

### 4.4 Predictor evaluation

All predictors were evaluated through per-protein cross-validation experiments. Unless otherwise noted, the training data for each predictor was first split into 10 randomly generated partitions, such that all data points from a given protein were in the same partition. Then, in an iterative manner, each partition was treated as the test set and the remaining nine partitions constituted the training set. To avoid information leak, feature selection, normalization and dimensionality reduction parameters were obtained on the training partition, and then applied to the test partition within each iteration. After 10 iterations, every data point was assigned a prediction score. These scores were then used to estimate the accuracy of the model.

Evaluation of MutPred2 was performed in a similar manner, except that the crossvalidation partitions were defined more stringently. Instead of a per-protein partition definition, a more stringent per-cluster partition definition was adopted as proposed by Calabrese *et al.* [30] Protein sequences in the data set were first clustered using CD-HIT at the 50% sequence identity threshold. We then ensured that all substitutions from the same cluster were either entirely in the training set or entirely in the test set.

### 4.5 Independent validation and comparison

For the purposes of additional evaluation and comparison with other methods, an independent test set was compiled from mutations deposited in ClinVar [31] (March 5, 2015) and UniProt [75] (“humsavar.txt”, release 2015 04). Fragments of length 25 residues centered at the mutation position were extracted and compared to similarly constructed fragments from the training sets of five methods (MutPred2, MutPred [10], PolyPhen-2 (both models), SNPs&GO [30] and FATHMM [26]) using CD-HIT-2D [76]. In the case of FATHMM, the additional “humsavar” data set that it was tested on (with similar performance) was used, because FATHMM training sets were not publicly available. All mutations whose local neighborhoods shared at least 50% sequence identity with those from at least one of the training sets were filtered out. Predictions for MutPred2, MutPred, and PolyPhen-2 were obtained using locally installed versions of the software. Scores for SNPs&GO were obtained through multiple queries to its webserver. The FATHMM scores, along with predictions for other methods such as CADD [25], SIFT [21], MutationTaster2 [28], GERP++ [27], and PhyloP [29] (20-way) were directly obtained from the dbNSFP database [77] (v3.0) of all possible single nucleotide substitutions. In cases where the chromosomal positions in this database mapped to multiple protein positions, one-to-one correspondence of isoforms (and positions) was verified. Although other methods for the prediction of pathogenicity exist, we chose this representative set based on recommendations recently made by the American College of Medical Genetics and Genomics and the Association for Molecular Pathology [11]. The entire procedure remained the same for the threshold of 80% sequence identity.

### 4.6 Score distributions on genomes

Two individuals from each of the five super-populations represented in the 1000 Genomes Project [34] (Phase 3) were randomly selected, such that they came from different populations. In total, variants for 10 genomes were extracted from the integrated variant call format (VCF) files: NA19026, HG02014, HG02002, HG01075, HG02384, NA18632, NA12829, HG01615, HG04206 and HG02651. ANNOVAR [78] was used to identify and retain nonsynonymous single nucleotide substitutions, map their coordinates to amino acid positions in protein sequences, extract their zygosity information and obtain minor allele frequencies (MAFs) from the ExAC browser [50]. The “coding change.pl” program in ANNOVAR was used to obtain protein sequences for MutPred2. Both MutPred2 and PolyPhen-2 (both models) were locally installed and run on this data set. The resulting scores were binned into fixed intervals for each individual separately. The mean fraction of variants within each bin and its standard error over all 10 individuals were then plotted. Similarly, in the case of MAFs, the mean allele frequency and its standard error were plotted.

Enrichment of properties in the MutPred2 disease set. Frequently altered properties in the set of all disease variants in MutPred2’s training set were identified by first deciding a threshold for loss and gain scores based on a predetermined FPR (here, 1%). Assuming that the vast majority of the non-pathogenic substitutions do not affect protein properties, one can use the fraction of these substitutions with a score greater than or equal to the threshold to approximate the rate. For instance, if an FPR of 5% is desired and if 5% of non-pathogenic variants have a loss score of 0.4 or greater, then the threshold would be 0.4. Based on this threshold, the numbers of disease-causing variants with and without the given mechanism were then counted. Thus, two proportion values were obtained: one for the fraction of disease-causing variants affecting the property (*d*_*f*_), and one for the fraction of non-pathogenic variants affecting the property (*n*_*f*_). Then, the enrichment *E* was calculated as

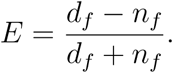

If *E* was positive, the property was considered to be enriched, and if it was negative, the property was treated as depleted in the disease set. Significance to these enrichment/depletion values was assigned using a one-sided Fisher’s exact test with subsequent correction for multiple-hypothesis testing, using the Benjamini-Hochberg method [79]. Since PTMs are known to occur on specific residues, these were further divided into two separate categories when generating the counts: when a substitution occurred at the predicted PTM site exactly, and when it occurred in its neighborhood. Note that although this data set was dominated by mutations from HGMD, the diseases covered are not strictly monogenic. Nevertheless, we refer to this set as the data set of Mendelian diseases.

### 4.7 Analyses on neuropsychiatric disorder mutations

A data set of 4,324 *de novo* mutations identified through whole exome or whole genome sequencing of the individuals diagnosed with autism spectrum disorder (ASD), intellectual disability, epileptic encephalopathy and schizophrenia [42, 43, 48, 49, 52, 54, 67, 69, 80–97], along with a control set of 1,316 *de novo* mutations from the healthy siblings [42, 43, 52, 69, 86, 90, 92, 93, 95, 98] was curated from the published literature. Genes with mutations shared by both case and control sets were removed from the analyses. Unlike the MutPred2 disease set, there was no prior knowledge of which mutations in the case and control sets were pathogenic or benign. Therefore, MutPred2 pathogenicity scores (at the 5% FPR threshold; score of 0.79) were used to divide each set into pathogenic and benign mutations. Only the mutations above this score threshold were considered for further analyses. To identify structural and functional signatures for each substitution, property scores were ranked in decreasing order. Then, the fractions of substitutions with a given property in the top three were compared between the case and control sets using a one-sided binomial test. The resulting P-values were then FDR-corrected using the Benjamini-Hochberg method.

### 4.8 Yeast two-hybrid assays

Candidate genes for experimental validation were selected based on their MutPred2 scores and the availability of the corresponding clones in the human ORFeome v8.1 collection [47]. The common SNPs in close proximity to the potential disease mutation were extracted from dbSNP. We also verified that the selected SNPs are not present in ClinVar. All genes are stored in ORFeome v8.1 in the pDONR223 vector.

Missense mutations were introduced into the ORFs by the site-directed mutagenesis using PCR overlap [99]. The M13 primers were used as the flanking primers for the PCR overlap reactions, and the sequences for the forward (5’ to 3’ direction) primers used for the sitedirected mutagenesis were as follows:

`STXBP1 R551C: GAGCCTGAATGAGATGTGCTGCGCCTACGAGGTG;`

`STXBP1 G544D: CATTTTCATCCTTGGGGATGTGAGCCTGAATGAG;`

`STXBP1 G561R: GTGACCCAGGCCAACAGAAAGTGGGAGGTG;`

`STXBP1 A113T: CTGACTCTTGTCCAGATACCCTGTTTAATGAACTG;`

`STXBP1 V84I: CATCCGAGAAGTCCATCCACTCTCTCATC;`

`ZBTB18 R486G: GTACAGCTCGGTGGTCTCGGAACTGGGCATCTCC;`

`ZBTB18 G416R: GTGCTCGCTGTGTAGGAAGACTTTCTC;`

`ZBTB18 A502T: GGTCAAAAGCGAAACACTGAGCTTGCC;`

`ZBTB18 T507A: CTGAGCTTGCCTGCTGTCAGAGACTG;`

DNMT3A R635W: GAGAAGAGGAAGCCCATCTGGGTGCTGTCTCTCTTTG.

The conditions for the PCR reaction were as follows: 94*°*C for 5 minutes, 30 cycles; 94*°*C for 30 seconds, 55*°*C for 30 seconds, 68*°*C for 1 minute in the PCRs 1 and 2 and for 2 minutes and 30 seconds in the PCR 3; and lastly 68*°*C for 7 minutes. The resulting mutant ORFs were Gateway cloned into the pDONR223 vector, and verified by Sanger sequencing. Then, the mutant ORFs were Gateway cloned into the pDB DEST vector and transformed into yeast for pairwise interactions testing. Briefly, miniprep plasmid DNA of all DB-X clones, both wt and mutant constructs, were transformed into the yeast two-hybrid strain MAT*α*Y8930. The interacting partners of the wt proteins were extracted from BioGRID [100]; only the partners that are present in the human ORFeome v8.1 collection were subsequently tested for interactions. The binary protein-protein interaction yeast two-hybrid (Y2H) screens of all DB-X baits against AD-Y preys (i.e., partners) were performed using previously described methods [64, 101]. Briefly, the DB-X and AD-Y clones were mated in YEPD media for 24h and then plated on Sc-Leu-3AT and Sc-Leu-His-CHX plates (i.e., test for autoactivation) for selection. Only colonies that grew on the Sc-Leu-3AT plate but not on the Sc-Leu-His-CHX plates were counted as positives. All the pairwise Y2H screens were repeated three times in independent experiments and only interactions that scored as positives at least twice were considered as positives.

## Acknowledgements

We thank Prof. Matthew W. Hahn for helpful comments. This work was supported by the US National Institutes of Health (NIH) grants R01 LM009722 (SDM), R01 MH105524 (LMI and PR), MH104766 (LMI), R01 MH076431 (JS) and the Indiana University Precision Health Initiative (PR). DNC and MM acknowledge Qiagen Inc. for their financial support through a License agreement with Cardiff University.

